# Deep learning for species identification of modern and fossil rodent molars

**DOI:** 10.1101/2020.08.20.259176

**Authors:** Vincent Miele, Gaspard Dussert, Thomas Cucchi, Sabrina Renaud

## Abstract

Reliable identification of species is a key step to assess biodiversity. In fossil and archaeological contexts, genetic identifications remain often difficult or even impossible and morphological criteria are the only window on past biodiversity. Methods of numerical taxonomy based on geometric morphometric provide reliable identifications at the specific and even intraspecific levels, but they remain relatively time consuming and require expertise on the group under study. Here, we explore an alternative based on computer vision and machine learning. The identification of three rodent species based on pictures of their molar tooth row constituted the case study. We focused on the first upper molar in order to transfer the model elaborated on modern, genetically identified specimens to isolated fossil teeth. A pipeline based on deep neural network automatically cropped the first molar from the pictures, and returned a prediction regarding species identification. The deep-learning approach performed equally good as geometric morphometrics and, provided an extensive reference dataset including fossil teeth, it was able to successfully identify teeth from an archaeological deposit that was not included in the training dataset. This is a proof-of-concept that such methods could allow fast and reliable identification of extensive amounts of fossil remains, often left unstudied in archaeological deposits for lack of time and expertise. Deep-learning methods may thus allow new insights on the biodiversity dynamics across the last 10.000 years, including the role of humans in extinction or recent evolution.

## Introduction

The identification of species has been a key issue in biology since Linnaeus (1758) and it remains a very important aspect for describing the past and extant biodiversity, with applications from conservation strategies (Amori et al. 2009), to the study of wildlife reservoirs of zoonoses (Müller et al. 2013). Molecular data have now widely replaced morphological criteria for such identification purposes (Kress and Erickson 2008; Vallejo and González-Cózatl 2012). Such methods even allow to identify species from degraded or environmental samples (Galan et al. 2012). However, even genetic identifications of species require to be based on properly identified specimens, including morphological aspects applicable to museum specimens (Müller et al. 2013). Furthermore, in the fossil record, morphological criteria are often the only data available for the description of the past biodiversity. In an archaeological framework, documenting the anthropogenic impact on vertebrate evolution since the Late Glacial period fostered the development of methods of numerical taxonomy to reach reliable identifications at the specific and intraspecific levels (Thomas Cucchi et al. 2014; Thomas Cucchi et al. 2017; Thomas Cucchi et al. 2020; Thomas Cucchi et al. 2019; Hulme-Beaman et al. 2018; F. James Rohlf and Marcus 1993; Stoetzel et al. 2017). Recent studies also explore morphological markers of behavioral change in order to document early steps of the domestication process (Harbers et al. 2020; Owen et al. 2014; Seetah et al. 2014) that has driven the evolutionary trajectory at the root of modern societies (Vigne 2015).

The optimization of species identification based on morphological criteria therefore remained a relevant field of research, to explore the fossil record, and to document extant biodiversity in countries where molecular studies remain difficult to perform. Many quantitative studies of morphological differences between species have been based on biometric measurements, especially on craniofacial characters [e.g. (Barčiová and Macholán 2009; Chassovnikarova and Markov 2007; M. E. Taylor and Matheson 1999)], an approach in which size differences are however very important. The rise of geometric morphometrics (GMM) (Adams et al. 2013; F. James Rohlf and Marcus 1993) provided efficient tools to further investigate morphological differences between species (Cordeiro-Estrela et al. 2008; Coster and Field 2015; Jaramillo-O et al. 2015; McGuire 2011). By allowing a separate analysis of size and shape, these methods notably allowed to disentangle these two important components of morphological evolution.

The current rise of new methods in machine learning for computer vision (Christin et al. 2019) raises the question of their pertinence to provide automated, reliable species identification based on morphology. Indeed, breakthrough performances have been achieved in the deep-learning era (Krizhevsky et al. 2012), in different contexts where a classification task has to be performed using images, such as for species identification (Wäldchen and Mäder 2018). The principle is simple: a deep neural network model has to be trained on a large set of labeled images (here, the label would indicate the species) so that the model can classify new images with a high predictive performance as well as a very high computing rate (dozens of images per second). If successful, such machine learning methods could constitute a fast and efficient alternative to GMM methods, which require time and expertise for successful applications. To investigate the potential of such approaches, and their transferability to the fossil record, the present study attempted to apply deep learning algorithms to the identification of three rodent species as case study, based on pictures of their molar tooth row. The performance of the deep learning identification was compared with the geometric morphometrics approach using.

### Case study: identification of three rodent species based on their molar morphology

Rodents represent the most diverse order of mammals, with ca. 2000 species including nearly half of the mammalian species (Wilson and Reeder 1993). They are pests for harvest and can be important reservoirs of zoonoses; in this context, species identification can be important for management (Heroldová and Tkadlec 2011; Müller et al. 2013). Recognizing even closely related species can be important for understanding their ecology and distribution in the landscape. As a consequence, efforts are still being done to elaborate criteria for species identification based on external morphology but also on craniofacial measurements, especially on skull and mandible (Barčiová and Macholán 2009; Javidkar et al. 2007; Pimsai et al. 2014; Siahsarvie and Darvish 2008; P. J. Taylor et al. 1995). Molar teeth bear important phylogenetic information in this group (Misonne 1969) and many identification keys integrate tooth measurements (Javidkar et al. 2007; Pimsai et al. 2014; Siahsarvie and Darvish 2008). Molar teeth are also very important because they constitute the most frequent fossil remains for such small mammals with fragile skulls. Their study thus provides irreplaceable insights into the biodiversity of former rodent faunas (López-Antoñanzas et al. 2019; J. Michaux 1983; Misonne 1969; P. J. Taylor et al. 1995). In the archaeological context, disentangling the house mouse from its wild counterparts delivered precious insights into the role of human niche construction in the emergence and spread of the commensal house mouse as well as the dynamics of human settlements (Thomas Cucchi et al. 2013; Thomas Cucchi et al. 2020; Thomas Cucchi et al. 2005; Weissbrod et al. 2017).

The three species considered in this case study are the house mouse *Mus musculus* (subspecies *domesticus*), the European wood mouse *Apodemus sylvaticus* and the Cairo spiny mouse *Acomys cahirinus*. All are murid rodents (Muridae family) but while the house mouse and the wood mouse are murine rodents (Murinae sub-family), the spiny mouse belongs to another sub-family, the Deomyinae (Steppan and Schenk 2017). However, *Acomys* displays an important morphological convergence, especially regarding tooth morphology, with murine rodents (Denys et al. 1992) and only molecular methods were able to evidence its attribution to another sub-family (Chevret et al. 1993).

These species are easily recognizable based on external morphology, and for specialists, even based on molar morphology (Fig. 1). They thus provide a case study of close but recognizable morphologies to test the relative performance between GMM and deep learning approaches. Furthermore, previous studies on these species (Renaud et al. 2020; Renaud et al. 2017; Renaud and Michaux 2003; Renaud et al. 2015) made available a high number of pictures to feed the deep learning approach with well-identified modern specimens. The test focused on the first upper molar (UM1) which bears most of the phylogenetically relevant characters in murine rodents (J. Michaux 1983; Misonne 1969). Focusing on this molar tooth only also allowed to test the transferability of this approach to fossil material, mostly composed of isolated molars.

**Figure 1.**
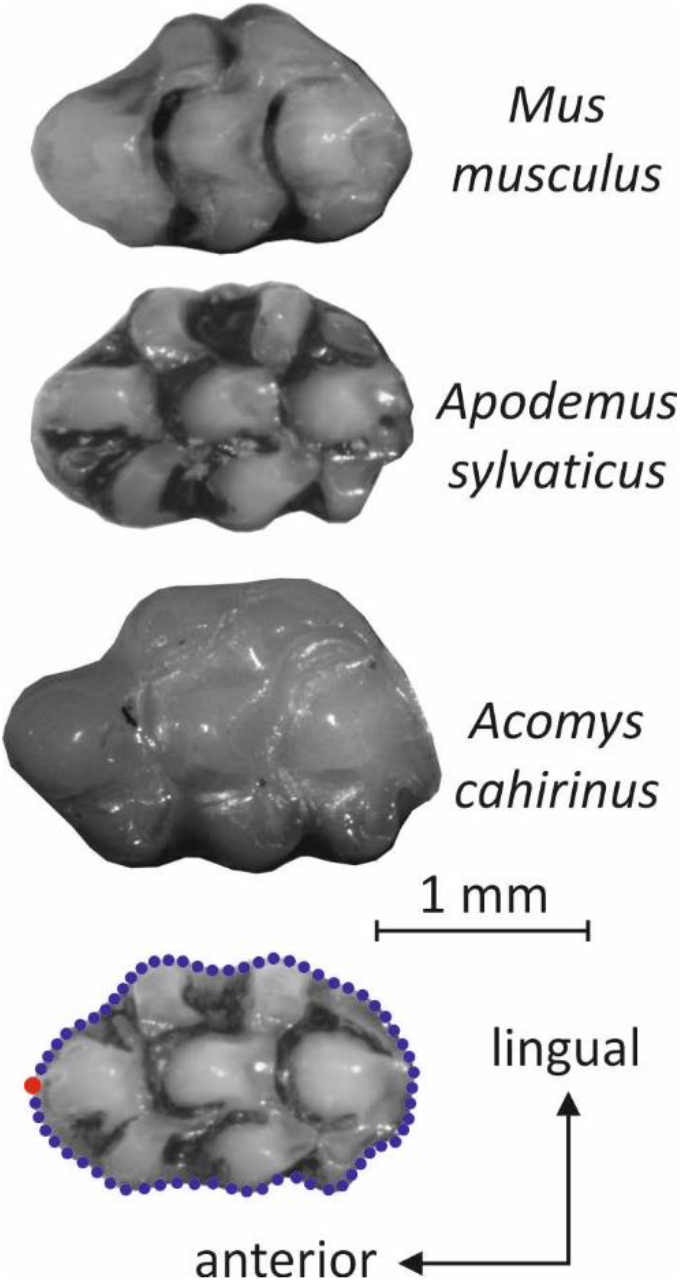
Pictures exemplifying the morphology of the first upper molar in the house mouse (*Mus musculus*), the European wood mouse (*Apodemus sylvaticus*) and the Cairo spiny mouse (*Acomys cahirinus*). All pictures to the same scale. The points delineating the occlusal outline and used for the GMM analysis are depicted on a wood mouse molar as blue dots (in red the starting point).

The objectives were thus the following: based on a sample of almost 1500 pictures of molar rows of modern animals, a protocol of automatic cropping of the UM1 followed by an automatic classification procedure was elaborated, both steps being based on deep learning. The deep learning classification efficiency was then compared with the results of a geometric morphometric analysis of the molar outline, computed on the same modern dataset. Second, the classification performance of both approaches was assessed on fossil molars. Finally, practical advices for the elaboration of a dataset to foster efficient deep learning approach were gathered from this trial procedure.

## Material and Methods

### Material for the modern referential (Supp. Table)

#### Spiny mouse (*Acomys cahirinus s.l*.)

This species was represented by 96 animals, documenting the morphological variation in the Eastern Mediterranean area. Twelve specimens from the Museum National d’Histoire Naturelle (Paris, France) documented the morphology of North African populations, including eight animals identified as *A. cahirinus* from Cairo, Egypt (vouchers: 2001-11; 1997-1308; 1996-432; 1996-446; 1996-431; 1996-430; 1994-1280; 1999-6); and four other specimens attributed with less certainty to *A. cahirinus*, coming from Sudan and Chad (vouchers: 1906-118a; 1906118b; 1906-118c; 1981-1059). This sampling was completed by specimens from Crete (61), Cyprus (6) and Turkey (17). The context of isolation on the islands of Crete and Cyprus, and to some extent in the restricted patch where *Acomys* is found in Turkey, drove morphological differences in tooth size and shape (Renaud et al. 2020).

#### Wood mouse (*Apodemus sylvaticus*)

This species is native to Europe and northwestern Africa. The dataset included 588 wood mice. A first set of 264 animals was previously used to characterize geographic patterns of differentiation related to latitude and insularity (Renaud and Michaux 2003, 2007). This set included wood mice from various places in continental Western Europe and North Africa: Germany (4), Switzerland (2), Belgium (19), France (86), Italy (40), Spain (43), Portugal (3), Bulgaria (2) and Tunisia (8). Specimens from different Mediterranean and Atlantic islands were further included: Oleron (15), Ré (7) and Yeu (1) in the Atlantic Ocean off Western France; Corsica (8), Elba (1), Ibiza (9), Sicily (15) and Marettimo (1) in the Western Mediterranean Sea. This sampling was completed by animals integrated in a study devoted to patterns of within-population variation of the first upper molar (Renaud et al. 2015). It included specimens from continental Italy (3) and France (205), as well as from the islands of Noirmoutier (5), Porquerolles (86) and Port-Cros (12), and Sardinia (13).

This sampling strategy covers the different phylogeographic lineages described so far (Herman et al. 2017; J. R. Michaux et al. 2003) as well as various insular populations. Almost all specimens have been genotyped, confirming their attribution to *Apodemus sylvaticus*. The specimens from the collection of the Museum National d’Histoire Naturelle (Paris, France) come from Western France, outside the distribution area of the morphologically close species *A. flavicollis*.

#### House mouse (*Mus musculus*)

This species is the typical commensal mouse associated with human settings; all the specimens in the dataset belong to the Western European subspecies *Mus musculus domesticus*. The sampling encompasses continental populations from France (224), Italy (40), Germany (14), Denmark (14) and Iran (10) (Renaud et al. 2019; Renaud et al. 2013; Renaud et al. 2017; Renaud et al. 2011) and various insular populations from Corsica (74) and Sardinia (11) (Renaud et al. 2011), from Orkney islands (82) and Madeira (182) (Ledevin et al. 2016), and from two sub-Antarctic islands: the small Guillou island, part of the Kerguelen Archipelago (20) (Renaud et al. 2013), and Marion island (92) (Ledevin et al. 2016).

### Fossil material

#### Pleistocene *Apodemus sylvaticus*

Fossil teeth of wood mice attributed to *A. sylvaticus* were collected in fossil deposits from South France (Mas Rambault, Le Lazaret, Orgnac) and Lyon surroundings (Vergranne, Arbignieu). The dating of these deposits ranged from the Early Pleistocene (Mas Rambault, ~1.3 Ma) to the Late Pleistocene (Orgnac 3, 35000 years) and even the Holocene (Arbignieu) (Aguilar et al. 2002; Jeannet 1981; Mein 1990). This dataset includes 38 teeth in total (Deschamps 2004; Renaud et al. 2005).

#### Le Mesnil Aubry, Iron age

Archaeological teeth from *Apodemus* (6) and *Mus* (5) were collected in the Iron Age deposit of Le Mesnil Aubry, near Paris (Guadagnin 1983). The macroscopic traits of the molar morphology and the dwelling context to attribute the *Mus* remains to *Mus musculus*.

#### The sequence of Tuda, Corsica

The Monte di Tuda cave is located in the Northern part of Corsica. It is filled with a 2 m thick natural deposit corresponding to a 2500 years of small mammal archeological record (Vigne and Valladas 1996). The European wood mouse *Apodemus sylvaticus* has been introduced to Corsica during the late Neolithic (6000-5000 years BP), whereas the house mouse *Mus musculus domesticus* appeared later in Corsica, after the Bronze Age (~2500 BP) (Vigne 1992). Both species are thus well documented in the record of the Tuda cave. The fossils considered here have been retrieved during a preliminary excavation in 1988 from the five most superficial layers. This sampling corresponds to 77 *Apodemus* and 133 *Mus* first upper molars.

### Data acquisition: pictures

For the modern specimens, the skull was positioned on a bead bed and manually oriented so that the occlusal surface of the right upper molar row matched at best the horizontal plane. When the right side was damaged, the left upper molar row was photographed and the picture was mirrored. All images were oriented with the anterior extremity of the molar row to the left, and the lingual side above (Figs. 1, 2).

**Figure 2.**
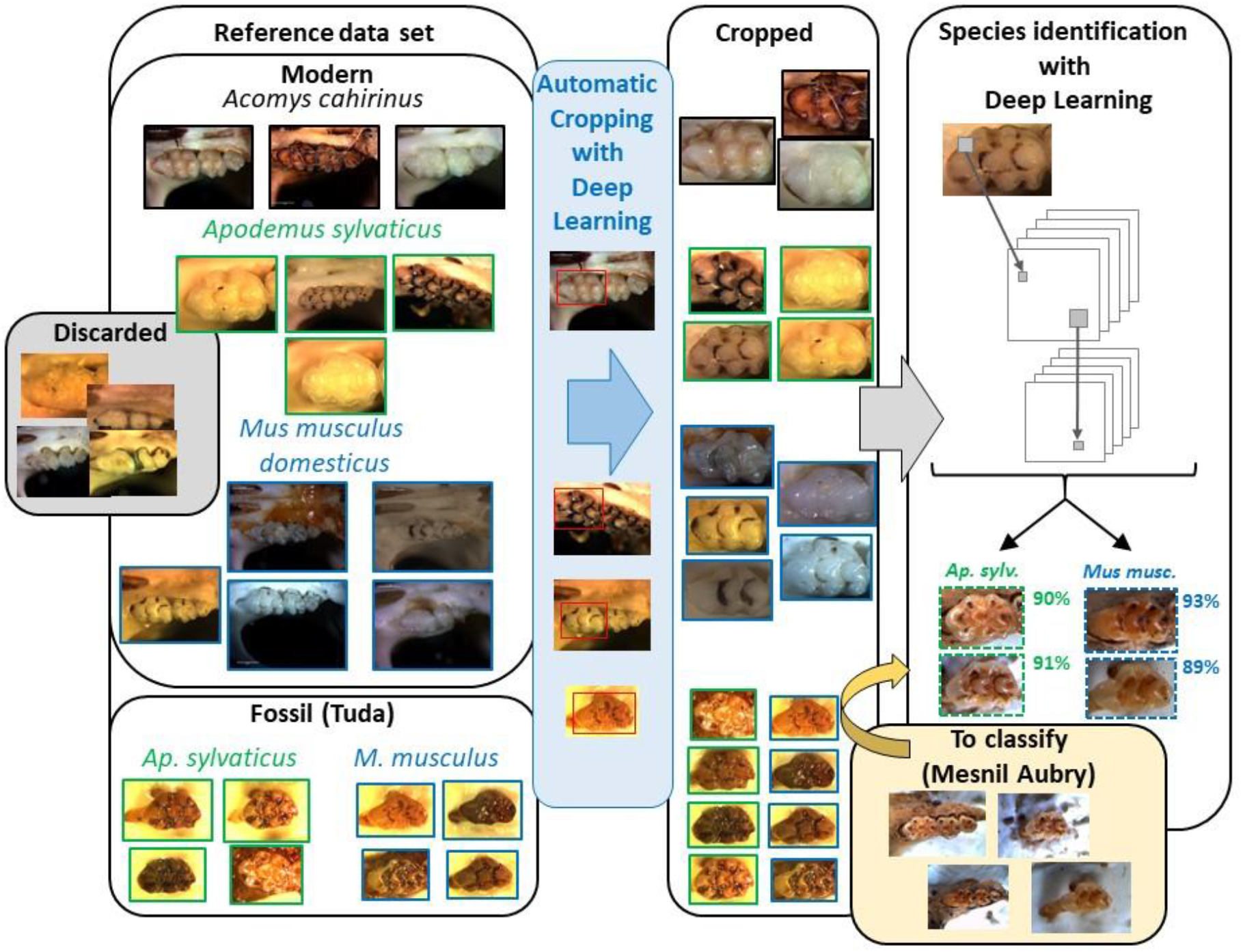
Schematic representation of the deep learning image-based procedure, from the initial images, to the cropped images and the classification results.

The pictures have all been taken using a binocular and a numerical camera. Lighting consisted of optical fibers that were manually adjusted to obtained a good picture. The pictures have been collected over more than 20 years, with different cameras and hence different resolution, magnification, and color balance.

Fossil teeth are most often found isolated. In this case, they are individually positioned with the occlusal surface up, the roots inserted in plasticine. Sometimes, the molars are still inserted in a broken jaw, as part of a fragmented molar row. The jaw is then positioned in plasticine. Pictures of the molars were taken using a binocular and a numerical camera in a similar way as used for modern specimens.

The same pictures were used for the geometric morphometric and deep learning approaches. For the later, few pictures were however discarded, because of molars too deeply worn down, or badly cleaned skulls hiding the relevant features on the teeth (Fig. 2; Supp Table).

### Methods for geometric morphometrics

The shape of the first upper molar (UM1) was described using 64 points sampled at equal curvilinear distance along the 2D outline of the occlusal surface using the Optimas software. The points along the outline were analysed as sliding semi-landmarks (Thomas Cucchi et al. 2013). Using this approach, the outline points are adjusted using a generalized Procrustes superimposition (GPA) standardizing size, position and orientation, while retaining the geometric relationships between specimens (F.J. Rohlf and Slice 1990). During the superimposition, semi-landmarks were allowed sliding along their tangent vectors until their positions minimized the shape difference between specimens, using the bending energy criterion. Because the first point was only defined on the basis of a maximum of curvature at the anterior-most part of the UM1, some slight offset might occur between specimens. The first point was therefore considered as a semi-landmark allowed to slide between the last and second points (Renaud et al. 2020). To compare the fossil UM1 to the modern dataset, all molars were superimposed in a same procedure.

The aligned coordinates were used as shape variables. The differentiation between the three species was analyzed using a leave-one-out cross-validated linear discriminant analysis (LDA). For the reclassification to the original groups, it removes one specimen at a time, and predicts its classification using LDA functions computed on all the remaining specimens. Classification accuracy is given by the percentage of specimens correctly assigned by the cross-validated LDA (cross-validated percentage, CVP). Fossil molars were considered as supplementary specimens reclassified to the groups of the modern dataset. The associated Canonical Variate analysis (CVA) computes axes maximizing the among-group relative to within-group variance; these axes can be used to visualize the differentiation between groups. Reclassified specimens can also be projected on these axes.

To assess how size differed between the three species and could influence identification based on dimensional data, the centroid size (CS) of the 64 points (i.e. square root of the sum of the squared distance from each point to the centroid of the configuration) was considered as an estimate of molar size.

The GPA was performed using the R package geomorph (Adams and Otarola-Castillo 2013). The Linear Discriminant Analysis with the cross-validated reclassification was performed using the package MASS (Venables and Ripley 2002). The computation of Canonical Variate Axes and the mean shapes for the different groups were obtained using the package Morpho (Schlager 2017).

### Methods for computer vision

#### Convolutionnal Neural Networks

The computer vision methodology used here relies on so-called “Convolutionnal Neural Networks” (CNNs) which are the backbone of many deep learning approaches (Krizhevsky et al. 2012). The key principle is the use of many sliding “neurones”, the “filters”, each dealing with a small set of pixels (for instance a 3×3 square) and sliding over the image to analyze every possible set of pixels. This operation is the “convolution” that is repeated many times with different filters. A CNN is a series of stacked “layers” containing a set of filters. The first “layer” slides on the image directly, whereas any of the following layers deals with the results of the previous layer. Therefore, these convolution layers are stacked one after the other, and connected, such that a CNN is a deep network of convolution layers. Different operators (e.g. “pooling”) are generally added to reduce the number of parameters and summarize the information all along the network. The convolutive part of the model, as explained here, is then followed by a classification part which can predict a value (when predicting a scalar) or score (when predicting a label). So, in summary, a CNN takes as input an image, process it with a series of convolutive operations, and finally returns a number which, in the present case, allows to predict a label (e.g. a species name). A CNN is therefore a model dedicated to image analysis, with a huge quantity of parameters that must be estimated using a large amount of data (here, images) available in a “training” dataset with labeled images. Since the parameters estimation procedure is iterative, we used as a starting point a pre-trained model (see details after), which already contained generic features that can be relevant to deal with our teeth images. This approach, called transfer learning (Shin et al. 2016), allowed us to deal with hundreds of annotated images only; otherwise, a CNN model must be trained with millions of annotated images.

#### Automatic molar cropping

An automatic cropping of the first molar (UM1) was performed in each image, using RetinaNet (Lin et al. 2017) which is a CNN-based tool for object detection (Zhao et al. 2019). RetinaNet is able to detect a range of object classes (e.g. a car, a person or an animal species) and displays the coordinates of a box around any detected object in an input image, plus a confidence score. The model has to be trained with annotated images, i.e. images for which the box coordinates are known. We used the pre-trained model with a ResNet50 backbone available along with RetinaNet. For a subset of images, we manually cropped bounding boxes around the first molars with the VGG Image Annotator (VIA) (http://www.robots.ox.ac.uk/~vgg/software/via/), obtaining 450 bounding boxes of modern UM1 and 88 boxes around fossil UM1 that were used as training data to enhance automatic molar cropping. Note that a large amount of manual bounding box delineation of images of fossil/archaeological rodent teeth was not required to train the model dedicated to fossil images, since we used the model previously trained to crop modern molars as a starting point, again using a transfer learning approach. Finally, the trained models were used to crop all the studied images, retaining the box with highest confidence score. For this step, we use the Keras implementation of RetinaNet available at https://github.com/fizyr/keras-retinanet.

#### Automatic species identification

A CNN-based method was designed to classify molar images into different classes, one class per species. We used the ResNet50V2 backbone available in Keras (https://keras.io/; input images resized to 300 x 300 pixels). Our CNN bakbone is followed by a softmax layer that gives a score for each class (all the score summing to 1), i.e. each species under consideration.

We evaluated four different CNN models, the first one being a 3-species model (identification of *Mus musculus, Apodemus sylvaticus* and *Acomys cahirinus*) whereas the others were 2-species models (identification of *Mus musculus* and *Apodemus sylvaticus*). The first model was trained with images of modern molars only. The second was trained with images of fossil molars only. The third was estimated using a transfer learning principle, using a model trained with modern molars as a starting point and then trained with images of fossil molars. The last model was trained using images of modern as well as fossil molars.

In all four cases, we adopted a 5-fold cross-validation procedure to compute the classification performance. We first randomly took out 20% of the original set of images to build a “validation” data set. The 80% remaining data constituted the “training” set on which we trained the model. We then evaluated the classification accuracy by predicting the species identification for all images of the validation data set, for which the actual species identification is known, and calculating the ratio between the number of well predicted images over the total number of images.

Finally, we estimated a fifth “reference” 2-species model using all the available data in the training set. Therefore, an evaluation of its accuracy was not possible, but presumably, its performance was improved by increasing the number of pictures used to feed the model. This model was used to classify molars from an archaeological deposit that was not included in the training model. This procedure was used as a test of “real case” application, when molars from a new deposit would have to be classified based on a preexisting reference dataset.

We trained each model with 10 epochs with batches of size 16. This pipeline was implemented with Keras 2.3.0.

## Results

### Differentiation between the species using the geometric morphometric approach

The three species are highly differentiated on the CVA axes (Fig. 3). The first axis (85.7% of between-group variance) opposes *Mus musculus* to *Apodemus sylvaticus*, whereas the second axis (14.3%) separates *Acomys cahirinus*. The general shape of the UM1 clearly differs in the three species. *Mus musculus* displays elongated UM1, with discrete posterior lingual cusps and a prominent anterior cusp. Molars are broader in *Acomys cahirinus*, but they share with *M. musculus* a relatively triangular posterior zone, and a well-delineated anterior cusp, especially on the lingual side. *Apodemus sylvaticus* also displays broad molars, but all posterior cusps are well expressed on the outline, and the anterior zone is short with a smooth transition with the next cusps (Figure 3A). The fossil specimens, projected on this space as supplementary specimens, felt within or close to the range of the corresponding species in the modern referential (Figure 3B).

**Figure 3.**
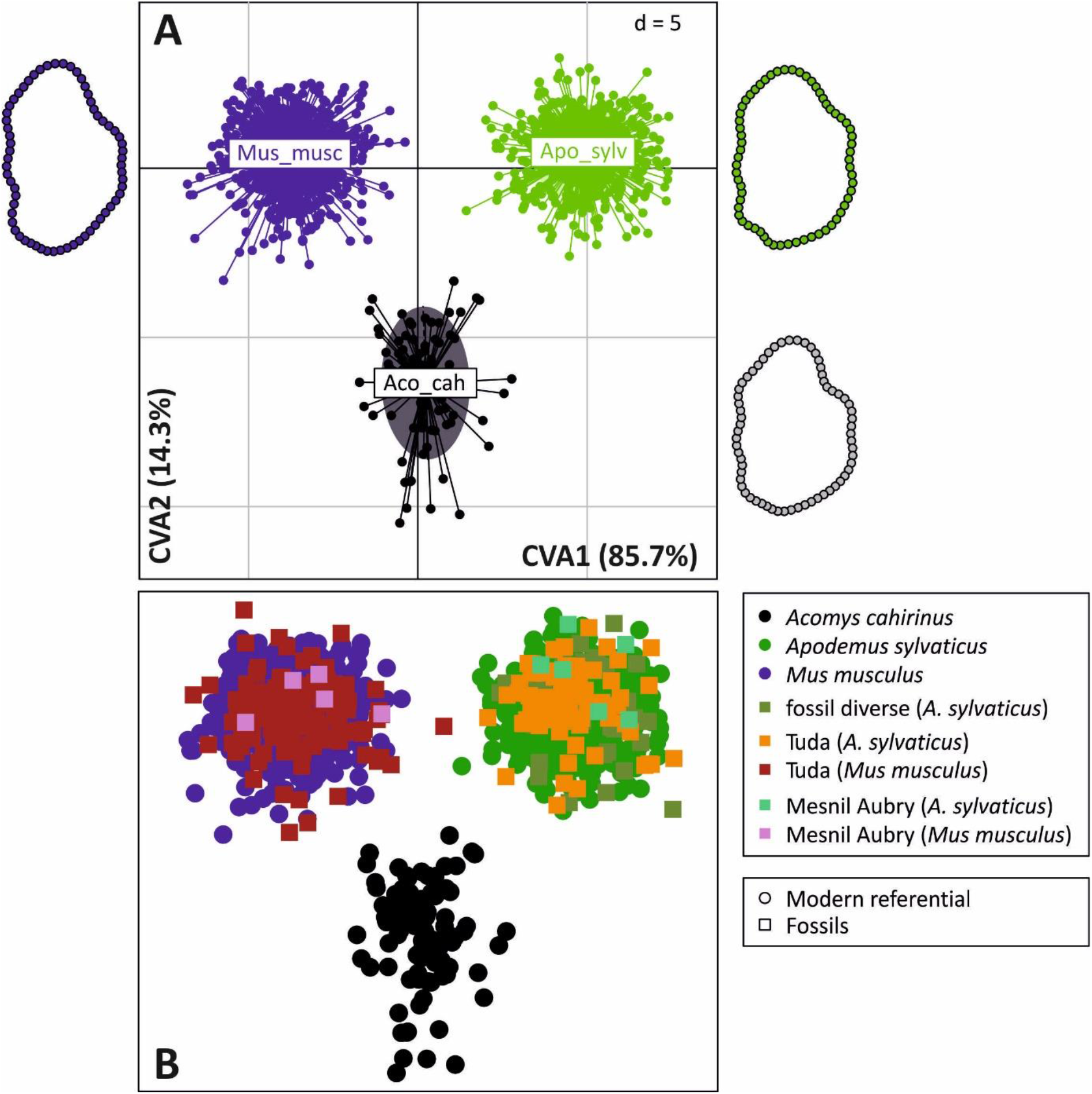
Morphological differentiation between the three species *Acomys cahirinus, Apodemus sylvaticus* and *Mus musculus* based on the geometric morphometric analysis of the first upper molar shape. A. Scores of the specimens on the first two axes of a CVA on the aligned coordinates. Close to the range of each species, a visualization of its mean molar shape. B. Projection of the fossil teeth as supplementary specimens on the same axes. MA: Mesnil Aubry.

The LDA on the aligned coordinates of the referential dataset provided 100% correct reclassification, even with the leave-one-out procedure (Table 1). Almost all fossils were also correctly reclassified, to the exception of one house mouse from Tuda.

**Table 1.** Reclassification of modern and fossil molars based on a geometric morphometric approach. Molar shape is described by the aligned coordinates of the points delineating the occlusal outline after a Procrustes superimposition. The modern referential was used as a referential in a Linear Discriminant Analysis and reclassified using a leave-one-out procedure. The fossil molars were considered as supplementary specimens and classified based on the discriminant axes computed on the modern referential.

|  |  |  | *Aco. cah.* | *Apo. sylv.* | *Mus. musc.* |
| --- | --- | --- | --- | --- | --- |
| Modern |  | *Aco. cah.* | 96 | 0 | 0 |
|  |  | *Apo. sylv.* | 0 | 588 | 0 |
|  |  | *Mus. musc.* | 0 | 0 | 763 |
| Fossils | Pleistocene | *Apo. sylv.* | 0 | 38 | 0 |
|  | Mesnil Aubry | *Apo. sylv.* | 0 | 6 | 0 |
|  |  | *Mus. musc.* | 0 | 0 | 5 |
|  | Tuda | *Apo. sylv.* | 0 | 77 | 0 |
|  |  | *Mus. musc.* | 0 | 1 | 132 |

Note that the three species differ notably in tooth size, *Mus musculus* displaying the smallest and *Acomys cahirinus* the largest molars (Fig. 4). Insular populations tended to display larger molars than their continental relatives. This insular variation increased the overlap between the different species. In some cases, the fossil teeth tended to be larger than their modern counterparts, for instance for the wood mouse and house mouse from Tuda, and the house mouse from Mesnil Aubry. Due to this variance in molar size, and to the high rate of correct classification based on shape only, including molar size in the predictors of the LDA (hence performed on log(Centroid Size) and the aligned coordinates) did not change the classification results.

**Figure 4.**
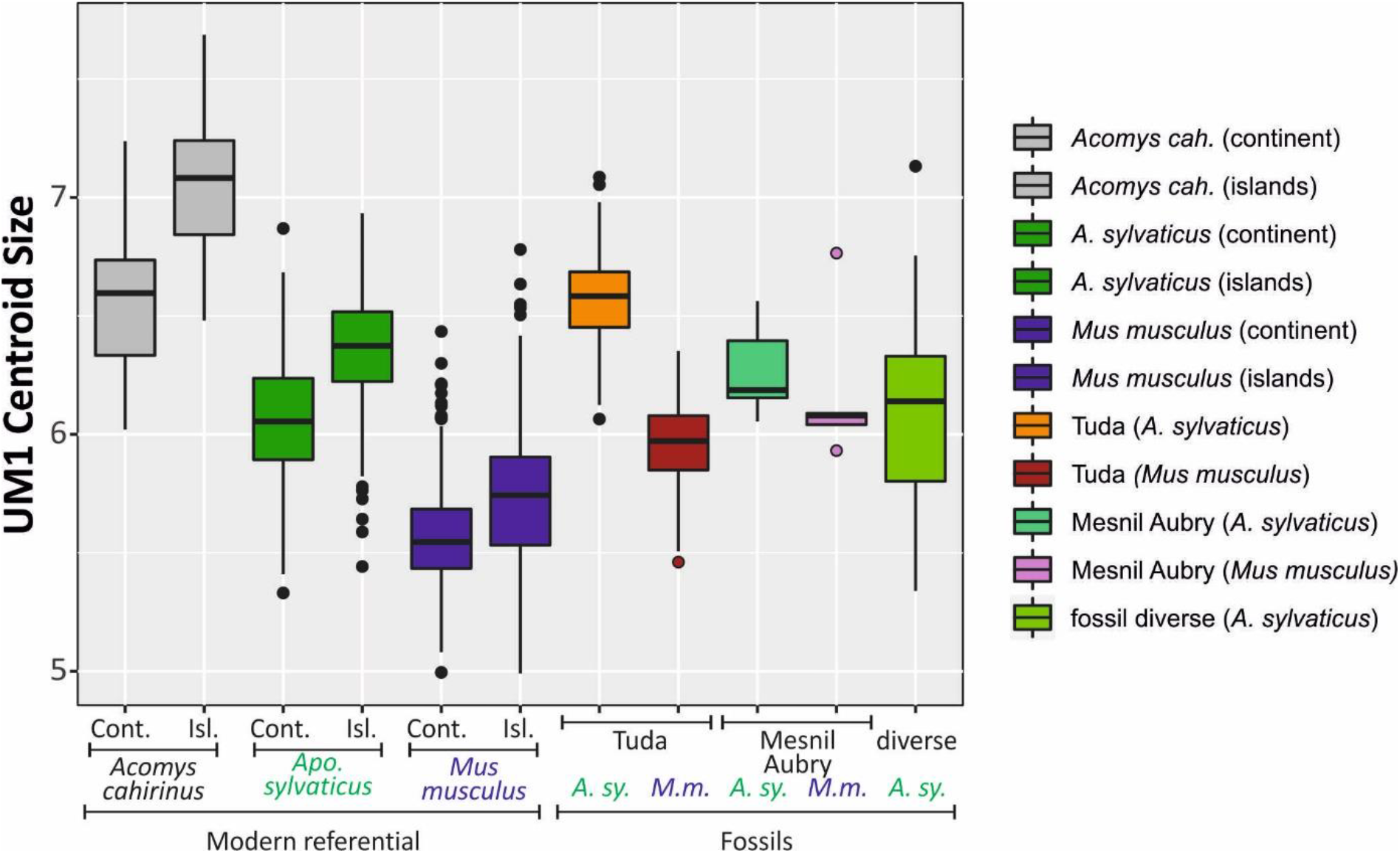
Differences in molar size between the three species, continental and insular populations, modern and fossil representatives. Molar size has been estimated by the centroid size of the points delineating the occlusal outline.

### Results for the computer vision approach

The UM1 was first cropped with the object detection method RetinaNet. From there, the predictive power of the deep learning procedure was evaluated for the four different settings (Fig. 5).

1. The CNN-based approach was very efficient in discriminating *Mus musculus, Apodemus sylvaticus and Acomys cahirinus* on modern UM1 images, with a large number of images in the training set (about 1200 images). The computed validation accuracies were close to 1. When focusing on *Mus musculus* and *Apodemus sylvaticus* modern molars, the performances were equally good (not shown).
2. Still focusing on *M. musculus* and *Apodemus sylvaticus* but dealing with a model trained on fossil molars only, poor accuracy performance (median < 90%) was obtained. This poor performance is due to the fact that the number of images in the training set was too restrained (about 150 images) to learn generic features that can be relevant for any image of fossil molar.
3. The next strategy was thus to involve transfer learning, relying on a model previously estimated on modern molars. This achieved a high accuracy performance in predicting the species for fossil molar images (accuracy > 98% in most cases).
4. An alternate approach consisted in pooling modern and fossil molar images in the training set. This approach raised good performance in accuracy prediction for both modern and fossil species. This last model was indeed able to capture some genericity that allowed for species identification in images of any of the two conditions (modern or fossil).

**Figure 5.**
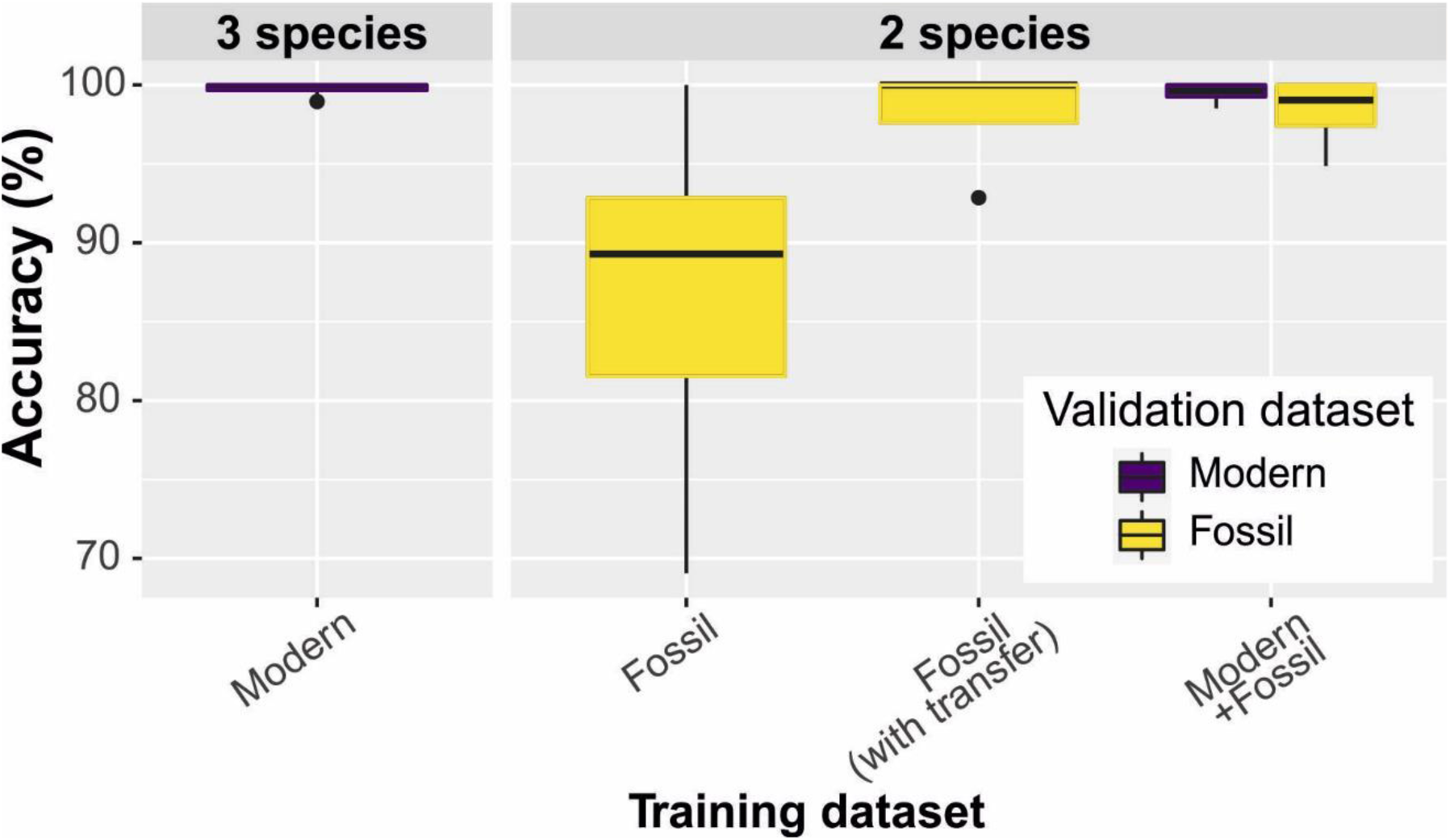
Accuracy of the deep learning models, computed in a repeated 5-fold cross-validation procedure. Four different CNN models were estimated with different training datasets (from left to right): 1) a 3-species model for modern molars, 2) a 2-species model for fossil molars without transfer learning; 3) a 2-species model for fossil molars with transfer learning; and 4) a 2-species for modern and fossil molars jointly. Each dataset was randomly split into 80% of the pictures used to train the model, and the remaining 20% used as a validation dataset. Accuracy values are computed on this 20% validation dataset.

The last step was to estimate a fifth « reference » 2-species model using all available data in the training set, therefore without letting 20% apart for the validation procedure. This reference model was use to classify the fossil molars from Mesnil Aubry (Fig. 6), that was never included in the training dataset, and thus constituted a “real case” of application to a new deposit. Except for one tooth identified as *Mus musculus* for which the model was unable to decide, the prediction based on the deep-learning procedure matched with good confidence (> 90%) the identification done by trained zooarchaeologists. Note that the pictures of Mesnil Aubry have been taken by another operator and with different camera and lighting settings from all pictures constituting the dataset used to train the model.

**Figure 6.**
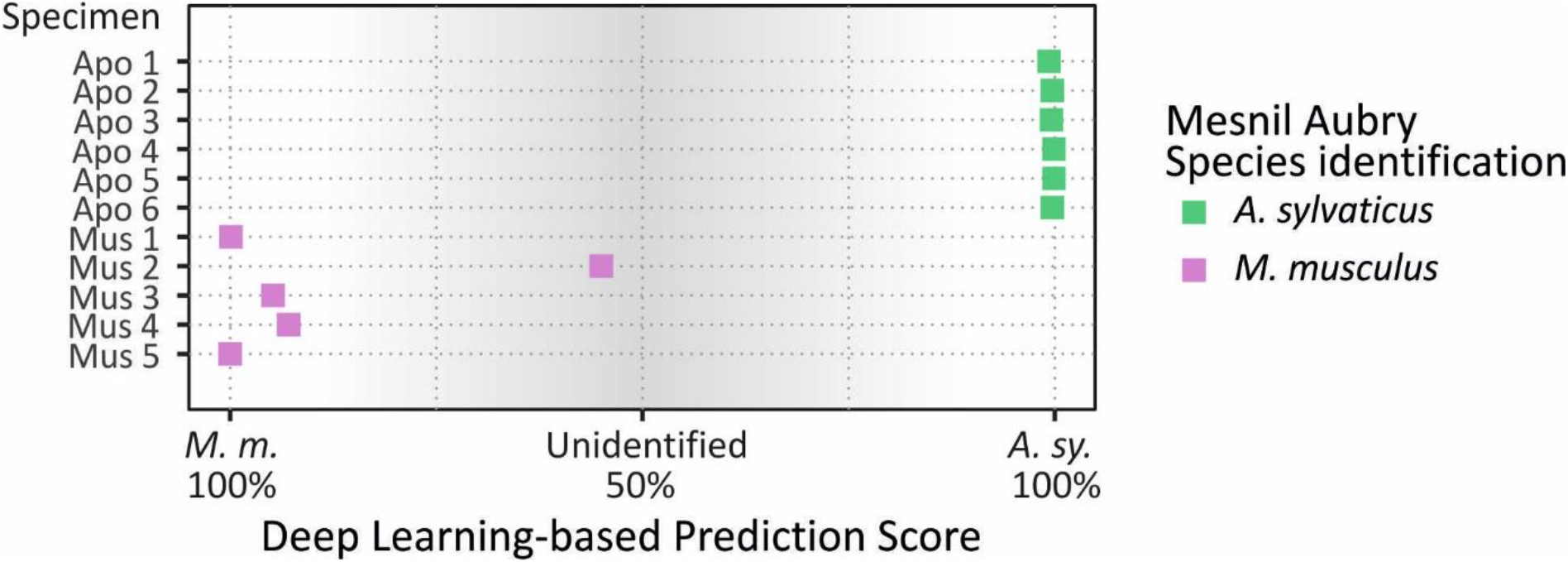
Deep learning-based classification of the fossil molars from Mesnil Aubry. A “reference” 2-species model was estimated with all the available images of modern and fossil molars, to the exception of those of Mesnil Aubry which were considered as an external dataset to be classified. The prediction score indicates to which species the specimen is the closest (both scores for predicting the two species sums to 100%), 50% indicating no decision. The grey zone therefore materializes scores for which the species prediction is not reliable. The prediction is compared to the actual species identification (color of the symbols).

## Discussion

The present study represents a proof-of-concept of the applicability of deep-learning algorithms to the identification of rodent species based on simple pictures of their molar rows, and of the transferability of this method to the identification of fossil teeth.

### Deep learning algorithms as a promising tool for the identification of species

In the case study presented here, the deep learning approach and the GMM analysis performed equally well, when the data set was large enough to adequately feed the deep learning models.

Geometric morphometrics became in the last decade the most performant method in numerical taxonomy, although biometric analyses remained common for zoological descriptions (Barčiová and Macholán 2009; Dianat et al. 2010; Javidkar et al. 2007; Pimsai et al. 2014). In most cases, GMM analyses require the manual acquisition of landmarks or outline data. In the case of the murine teeth, each molar occlusal view was manually delineated for the collection of the points along the outline, following a procedure applicable to different rodent taxa (Gómez Cano et al. 2013; Ledevin et al. 2010; Renaud et al. 1996). The data acquisition for the ~1500 mice presented here therefore required, in total, several weeks of full-time fastidious work of an experienced operator. The integration of datasets collected by different operators require a procedure of cross-measurement to check for inter-operability.

The deep-learning procedure requires the acquisition of a collection of well-identified pictures, like for the GMM analyses. However, once the pipeline is elaborated, our dataset only required few minutes to enter the set of thousand pictures and train the model, and only a few seconds to obtain classification results on few hundreds of pictures, using a graphics processing unit (GPU). Furthermore, running an existing pipeline of deep learning demands no in-depth knowledge of the biological group under study. In both case, the validity of the results is however fully dependent of the initial identification of the specimens composing the dataset used in the training step.

The interest of both methods is that this reference dataset can be elaborated from genetically-identified specimens, before a transfer to modern specimens without access to genetic analyses, or to fossil specimens. Importantly, both methods worked here without taking size into account. In the case of important size variations, for instance on islands (Millien 2006) or between fossil and modern representatives of the same species (Cassaing et al. 2011; Thomas Cucchi et al. 2014), this may be a crucial aspect for reliable classification results. Both methods will however face the limitation of transfer functions from the recent to the past. Going back in the past, the morphological divergence between the modern referential and the fossil sample may become too important for pertinent identifications. The case of the early Pleistocene wood mice from Mas Rambault however shows a resilience over one million year of evolution, but wood mice display a rather conservative morphological evolution (Renaud et al. 2005).

### Requirements for an efficient training dataset

The deep learning procedure learns relevant informations from image patterns. It is therefore important that the pictures of the reference dataset are not too different in their setting from the ones to be classified (Beery et al. 2018). A way to mitigate this issue and make the reference dataset transferable to a wide range of data is to make it as diverse as possible. This was achieved in our final “reference” CNN model by including an extensive sampling of modern and fossil molars. As a result, the model was able to almost completely classify pictures from Mesnil Aubry, which were taken by a different operator and with different settings than the pictures included in the reference dataset.

Achieving this diversity in the reference dataset should be done first, by including as much intraspecific variation as possible. For instance, in our case study, all the three species display important geographic variation, especially on islands (Ledevin et al. 2016; Renaud et al. 2020; Renaud and Michaux 2007). If the specimens to be reclassified come from insular environment, such as those from the Tuda sequence, it is advisable that the range of morphological variation encountered on different islands is included in the reference dataset. A too stringent cleaning of the reference dataset from damaged or aged specimens, with used teeth, may be in that respect counterproductive. Second, variation in the pictures themselves (lighting, color range, focus…) should ideally be included as well, to make the algorithm more resilient when facing a diversity of picture settings, and hence, more easily transferable to another set of pictures, possibly taken by other operators with other devices and conditions.

### Transferability to the fossil record

The possibility to classify fossil teeth with reference to genetically-identified modern specimens provide a great opportunity to document past diversity from the fossil record. However, the taphonomic processes during fossilization often alter the biological object, making a comparison of pictures taken on modern and fossil object not as straightforward. In the case of rodent fossil molars, a further issue is that data collected on modern specimens provide pictures of complete molar rows, with the first molar in contact with the second one, with the skull as background. In contrast, fossil teeth are found isolated and have to be inserted on plasticine to be properly oriented. The background of the first upper molar is thus radically different for modern and fossil specimens. Additionally, the enamel of fossil teeth often takes a different coloration from the original white tint. A mere transfer of the modern reference dataset to the classification of fossil teeth could thus appear problematic. A first way to deal with these issues was to develop an automated protocol to crop the UM1 from its background. A further way to mitigate this issue was to include some well-identified fossil specimens to the reference dataset, to make it more resilient to the background of the tooth.

### Potential ranges of applications

The potential of deep learning approaches for the identification of species has already been recognized, mostly for applications to the recognition of photos of living wildlife animals (Tabak et al. 2019; Wäldchen and Mäder 2018). When fed with enough pictures, such deep learning image-based methods can even achieve individual identification, as in the case of giraffes (Miele et al. 2020). The originality of the present study is however to explore applications dealing with pictures of biological material prepared for osteological collections, or even focusing on a single molar tooth as in our case study. In such context, providing fast and efficient identification tools, without the time and expertise invested in GMM methods, may be beneficial to reconsider ancient collections of museum specimens. For such purposes, the reference dataset could rely on pictures of the whole skull, which probably contain more phylogenetic information that the sole first upper molar (Rychlik et al. 2006).

Another important potential range of application for such algorithms regards the identification of fossil specimens, based on well-identified modern specimens. Despite limits due to inherent differences in pictures of modern and fossil specimens, our pilot study is highly promising in that respect. In the archaeological record, disentangling wild from domestic forms emerged as crucial to understand the process of domestication. GMM analyses allowed impressive progress in that respect (Balasse et al. 2016; Thomas Cucchi et al. 2016; Thomas Cucchi et al. 2009; Evin et al. 2013; Owen et al. 2014). Having recognized the importance of disentangling these close taxa in archaeological contexts, the application of fast deep learning strategies may widen the scope of such discriminations, reserving the use of sophisticated GMM analyses to an in-depth understanding of the signature of domestication on the different species and bones (Harbers et al. 2020).

A drawback of the deep learning strategy is that there is no or little feedback on the morphological characteristics allowing the discrimination of the taxa, precluding its use for enriching traditional determination keys. Indeed, CNNs are often used as black boxes (Wearn et al. 2019) and interpreting their parameters (i.e. having any idea of which image patterns were determinant for the classification) is still a research question (Miao et al. 2019; Selvaraju et al. 2017).

However, the potential for fast and performant specific identification could allow to deal with the extensive amount of fossil remains present in archaeological deposits, that are, for the time being, often left unstudied. These remains are very diverse and can include remains of small vertebrates but also insects, bivalves, crustaceans, etc. Provided that an extensive referential set of well-identified images can be elaborated, the deep-learning based identification of these remains may shed new light on the biodiversity dynamics across the last 10.000 years, including the role of humans in extinction or recent evolution.

This may allow to concentrate the application of GMM methods to studies devoted to the dynamics and processes driving morphological evolution: influence of the phylogenetic signature in interspecific evolution (Cardini 2003; Dianat et al. 2017), morphological convergence related to habitats and diets (Gomes Rodrigues et al. 2016; Samuels and Van Valkenburgh 2009), deciphering patterns of intraspecific variation (Thomas Cucchi et al. 2014; Monteiro et al. 2003; Renaud and Michaux 2007) up to characterizing patterns of covariation (Jamniczky and Hallgrímsson 2009) and the signature of allometry and developmental constraints on shape (Ferreira-Cardoso et al. 2020; Renaud et al. 2011).

The performance of deep learning systems in the field of zoology and archaeology will depend on the elaboration of extensive datasets of well-identified specimens, in order to train the models with as many pictures as possible. Using deep learning algorithms may democratize automated identification tasks, since writing only a few dozens of code lines can be sufficient to build a complete pipeline. However, despite being promising, these methods need to be rigorously evaluated to understand their potential limits and biases before extensive applications (Wearn et al. 2019). With adequate sampling, these methods could even deliver relevant results in the cases of sibling species, that remain sometimes difficult to disentangle using GMM methods (Dobigny et al. 2002).

## Acknowledments

We are grateful to all people who provided access to the biological material included in this study, especially Jean-Christophe Auffray, Jean-Pierre Quéré, Johan R. Michaux, Guila Ganem, Maria da Luz Mathias, Jean-Louis Chapuis, Benoit Pisanu, Frank Chan, George Mitsainas, Eleftherios Hadjisterkotis, Petros Lymberakis, and Ferhat Matur. We are further indebted to Pierre Mein and Jean-Denis Vigne for access to their collection of fossil material. We acknowledge Laure Deschamps for pictures of fossil material taken during a research internship. This manuscript is also a tribute to the late Anne Tresset, who provided the pictures of the material from Mesnil Aubry.

This work was performed using the computing facilities of the CC LBBE/PRABI. It was supported by funding from the French National Center for Scientific Research (CNRS) and the Statistical Ecology Research Group (EcoStat) of the CNRS.

---

**Supplementary Table.**
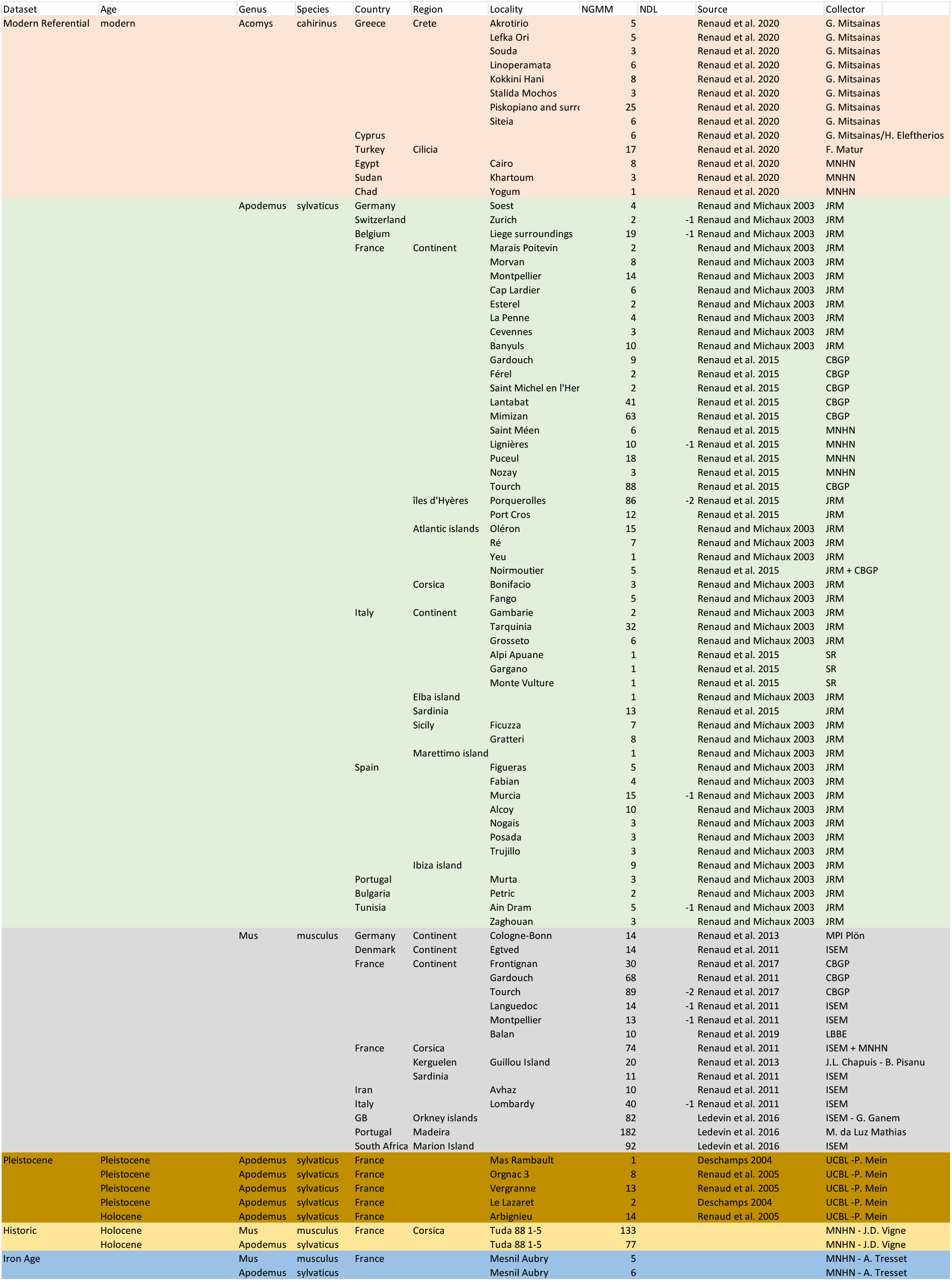

